# High-intensity cardiovascular exercise facilitates online motor skill learning, with no effect of BDNF genotype

**DOI:** 10.1101/2025.07.21.666051

**Authors:** Emily Brooks, Ziarih Hawi, Juliet Hosler, Jessica Kwee, Nermin Aljehany, Winston Byblow, Joshua Hendrikse, James Coxon

## Abstract

Previous studies have demonstrated that exercise can influence motor skill learning. However, the specific components of learning primed by exercise remain unclear. This study examined the effect of a preceding bout of high intensity interval training (HIIT) on the acquisition of a novel motor skill. The investigation focused on whether improvement in skill across the session was attributable to online gains during active practice or offline rest periods between practice blocks. There was also exploration as to whether common polymorphisms of the BDNF and DRD2/ANKK1 genes that regulate plasticity, learning, and memory, influenced the relationship between exercise and motor learning. It was demonstrated that HIIT enhanced skill acquisition, but that the effects of HIIT priming were not specifically attributable to within-session online or offline learning processes. Contrary to research on overnight consolidation, there was no interaction between BDNF, nor DRD2/ANKK1 genotype, with exercise primed skill learning.

## Introduction

Several converging lines of evidence have established a beneficial role of exercise in facilitating memory and learning^1–4^. Specifically, coupling motor task practice with acute exercise can promote improved skill learning, although the magnitude of the effect varies across studies^4^. What remains unclear is how exercise interacts with components of initial skill acquisition. Additionally, the variability in outcomes may in part reflect individual characteristics such as genetics^5^. Understanding the factors influencing this variability is important if we are to design paradigms to optimise the response to exercise, and maximise skill learning, as this may have practical implications for enhancing learning in clinical contexts.

Learning, and the formation of a motor memory, occurs dynamically over time through a combination of encoding/acquisition and consolidation processes. When learning new motor skills, acquisition describes the online manipulation and processing of new information during active practice, while consolidation processes are predominantly ‘offline’ during rest, strengthening new memories for retention and retrieval^6,7^. The most consistent finding in the literature to date is a priming influence of cardiovascular exercise on motor skill consolidation. Engaging in a bout of high intensity interval training (HIIT) paired with motor skill practice improves skill retention in the hours to days that follow^3,4,8–16^. However, the effect of cardiovascular exercise on motor learning during the acquisition phase remains comparatively unclear. Some studies have reported a faciliatory effect of exercise on motor skill acquisition^8,12,17^, though others have not demonstrated a benefit at this stage of learning^9,10,13,18,19^. For example, no effect of HIIT priming was observed for acquisition of an implicit serial reaction time task (SRTT)^19^. However, factors relating to experimental design may have limited the ability to detect effects in that study. Firstly, the faciliatory physiological effects and modulations in cortical excitability are known to be particularly prominent between 15-30 minutes following HIIT^20–24^, with closer temporal coupling of exercise and task practice providing greater benefits to skill learning^11,22^. However in the previous implicit study utilising the SRTT, practice took place on average 35 minutes post HIIT^19^, and therefore may not have been in the optimal time window to see a benefit on skill learning. Additionally, the SRTT is associated with a steep learning trajectory, with performance reaching a plateau after ∼4-5 minutes of practice^19,25,26^. Therefore, the absence of a HIIT-related influence may be attributed to the nature of the SRTT and the speed at which skill level plateaus, whereby the benefit of exercise may be masked, with both exercise and control groups quickly reaching an asymptotic skill level. It is plausible that for skills with different task demands (e.g., target selection, continuous visuomotor control, and force modulation) which have a more protracted learning trajectory to the SRTT, a benefit of exercise may be observed. Tasks like the sequential visual isometric pinch task (SVIPT) have a learning trajectory with a more prolonged ‘fast’ learning phase with further improvements across several days of practice^27^. Therefore, it is possible that using a task with a slower learning trajectory, and more optimally timed coupling of HIIT and task practice, HIIT may facilitate skill acquisition.

To examine the effects of HIIT on skill acquisition in a more granular manner, we sought to partition the session into online practice and offline rest periods. In recent years evidence has emerged of rapid offline consolidation of novel motor skills across a scale of seconds, during brief rest periods (e.g., 10s) separating blocks of active practice on motor sequence learning paradigms^25,26^. As exercise has been shown to particularly enhance offline consolidation^3,4,8–16^, it is possible that HIIT may enhance acquisition via improved offline consolidation over the practice session. In the aforementioned previous work from our group, we observed evidence of rapid offline consolidation of skill, but no additional benefit of HIIT exercise^19^, although this may be attributed to the aforementioned limitations of task timing and nature. Therefore, it remains unclear how HIIT influences the online and offline periods within a practice session to influence skill acquisition.

The impact of exercise on learning/memory processes varies across individuals^28^. Indeed, there is a growing body of evidence indicating that genetic factors, such as polymorphisms in the brain derived neurotrophic growth factor (BDNF) (Val66Met)^29–31^ and dopamine D2 receptor (DRD2/ANKK1 glu713lys) gene^5^ influence these processes. Studies have established that Val66Met is associated with reduced activity dependant secretion of BDNF^29–31^ and reduced memory function^31^. Additionally, the D2 receptor is critical for skill acquisition^32,33^. One study reported evidence that lys alleles in the DRD2/ANKK1 gene attenuate the benefit of exercise on skill learning, as measured across an overnight retention phase, with data pooled across several studies using different task designs^34^. This effect is yet to be reproduced with a prospective research design with a consistent task paradigm, and it is unclear how these polymorphisms influence interactions between exercise and motor learning during skill acquisition within a practice session.

Overall, the effects of HIIT exercise on motor skill learning during acquisition remain unclear. Similarly, whether these interactions occur online during active practice, or occur offline during brief rest periods is uncertain. Therefore, the aims of this study were to investigate the effect of a preceding bout of HIIT exercise on the acquisition of a novel motor skill. Individuals engaged in either a 20-minute bout of HIIT exercise (n = 27), or an active control, low intensity exercise (n = 24), prior to practicing a SVIPT task which was structured into alternating blocks of practice (online) and rest (offline). Three hypotheses were tested. First, greater skill learning would occur after HIIT exercise compared to active controls. Second, skills gains would manifest primarily during brief periods of rest (offline) than during practice (online). Finally, the benefit of HIIT exercise on skill learning would be less for individuals with BDNF gene and D2 receptor polymorphisms than those without.

## Results

### Participant Characteristics

HIIT and active control groups were well-matched for biological sex, age, BMI, level of physical activity, sleep quality, and genotype (Table 1). There was a 9 Newton (918 grams of force) difference in maximum voluntary pinch contraction (MVC) (BF_01_ = 0.40). A summary of the exercise parameters for the HIIT and active control sessions are provided in Table 2, and visualisation of percentage of heart rate reserve (%HRR = 100*((Exercise Heart Rate – Resting Heart Rate) / (Age-predicted maximal heart rate – Resting Heart Rate))) over sessions is represented in Figure 1a.

**Figure 1.**
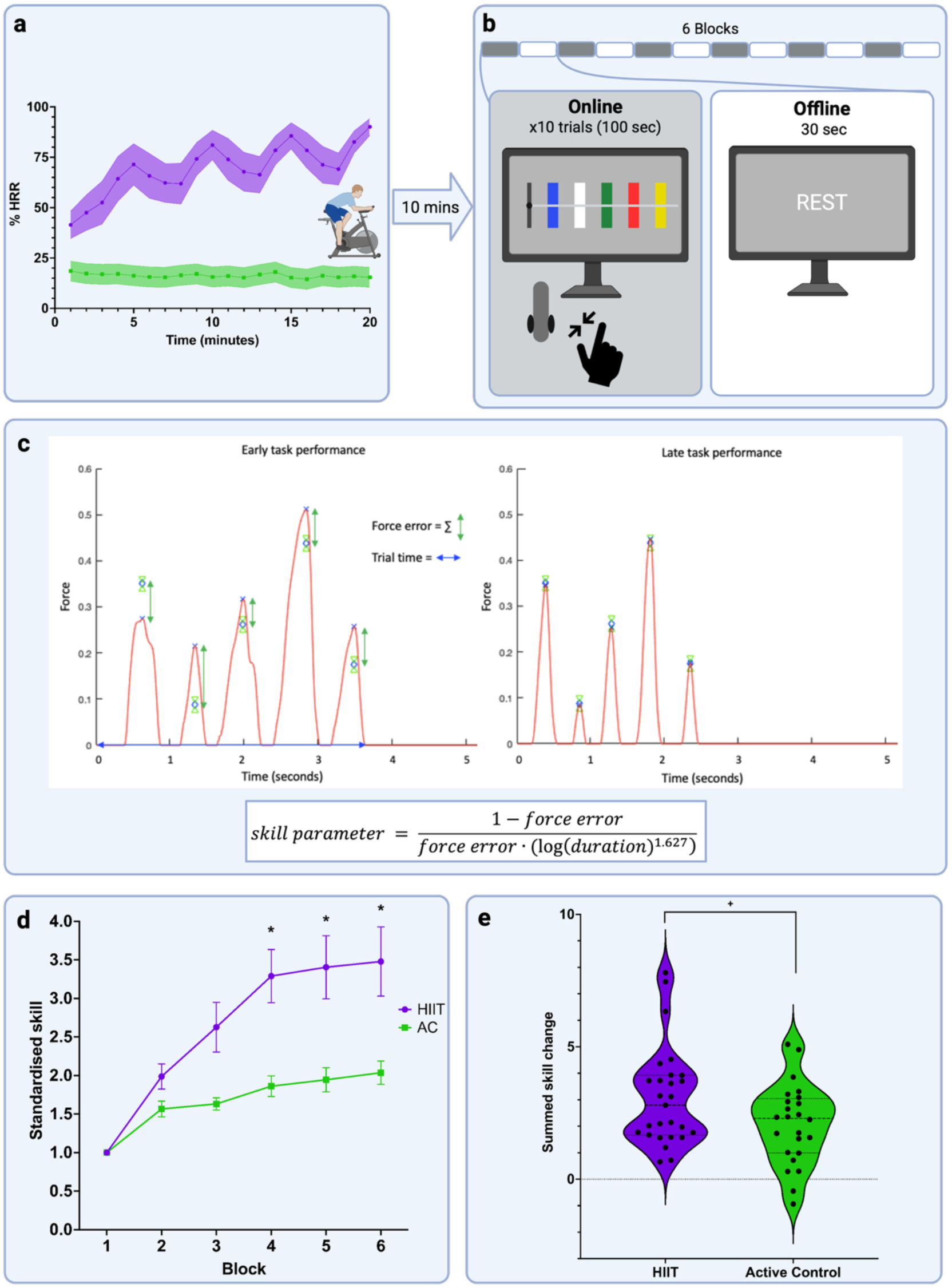
Paradigm overview of exercise and task, conceptualisation of skill, and skill learning results. *Note. **a)*** Overview of exercise and control protocols, represented as a percentage of heart rate reserve (HRR) as a measure of physiological workload, across the 20 minutes duration. Resting heart rate is 0 %HRR and maximum heart rate is 100 %HRR. The HIIT condition (purple) alternated between three minutes at around 50-60% HRR and two minutes of up to 90% HRR, while the active control (green) HHR remained stable at around 20%. **b)** Following ten minutes of rest after exercise participants completed the sequential visual isometric pinch task (SVIPT). The task was partitioned into online periods of 6 blocks of 10 trials, and offline rest periods of 30 seconds between blocks. **c)** Examples of individual SVIPT trials in early and later stages of practice. SVIPT skill is derived from speed (time taken to complete trial) and accuracy (summed difference between force peak and target). **d)** SVIPT skill over the session, quantified as a standardised score relative to baseline performance. Group differences are apparent from Block 2 of motor practice, with a significant difference observed from block 4 onwards, as indicated by the asterisks, * *all p* < .036. **e)** HIIT enhanced total learning on the SVIPT compared to the active control group ^+^(*p* = .043).

**Table 1.** Baseline Participant Demographics for the HIIT and Active Control Groups (Mean ± Standard Deviation).

|  | HIIT | Active control | Test statistic |
| --- | --- | --- | --- |
| N (% Female) | 27 (51.85%) | 24 (62.5%) | $\chi^2(1) = 0.96, p = 0.33$ |
| Age | 21.78 ± 2.21 | 22.33 ± 4.42 | $t(49) = 0.58, BF_{01} = 3.10$ |
| Body Mass Index (BMI) | 22.48 ± 3.37 | 23.2 ± 4.23 | $t(49) = 0.68, BF_{01} = 2.95$ |
| Physical Activity (IPAQ) | 3594 ± 1970 | 3995 ± 2902 | $t(49) = 0.59, BF_{01} = 3.09$ |
| Sleep quality (PSQI) | 4.25 ± 2.15 | 5.13 ± 2.77 | $t(49) = 1.25, BF_{01} = 1.89$ |
| Maximum voluntary contraction (MVC) | 54.12 ± 15.06 | 44.93 ± 21.38 | $t(49) = -2.34, BF_{01} = 0.40$ |
| BDNF Val/Val:Met carriers (Met/Met) | 9:15(4) | 12:11(0) | $\chi^2(1) = 0.53, p = 0.47$ |
| DRD2/ANKK1 Glu/Glu:Lys carriers (Lys/Lys) | 13:13(2) | 11:12(3) | $\chi^2(1) = 0.02, p = 0.89$ |

*Note.* Age is in years. Body Mass Index (BMI) was calculated as each participant's weight divided by their height squared ( $\text{kg/m}^2$ ). International Physical Activity Questionnaire (IPAQ) values are in metabolic equivalent units (Mets), indicating general level of physical activity. Pittsburgh Sleep Quality Index (PSQI) was the global score, indicating overall sleep quality. Maximum Voluntary Contraction (MVC) is reported in Newtons. BDNF shows the number of Val/Val homozygotes compared to the number of Met carriers (Val/Met and Met/Met) with the number of Met/Met homozygotes in brackets for each group. DRD2/ANKK1 shows the number of Glu/Glu homozygotes compared to the number of Lys carriers (Glu/Lys and Lys/Lys) with the number of Lys/Lys homozygotes in brackets for each group.

**Table 2.**
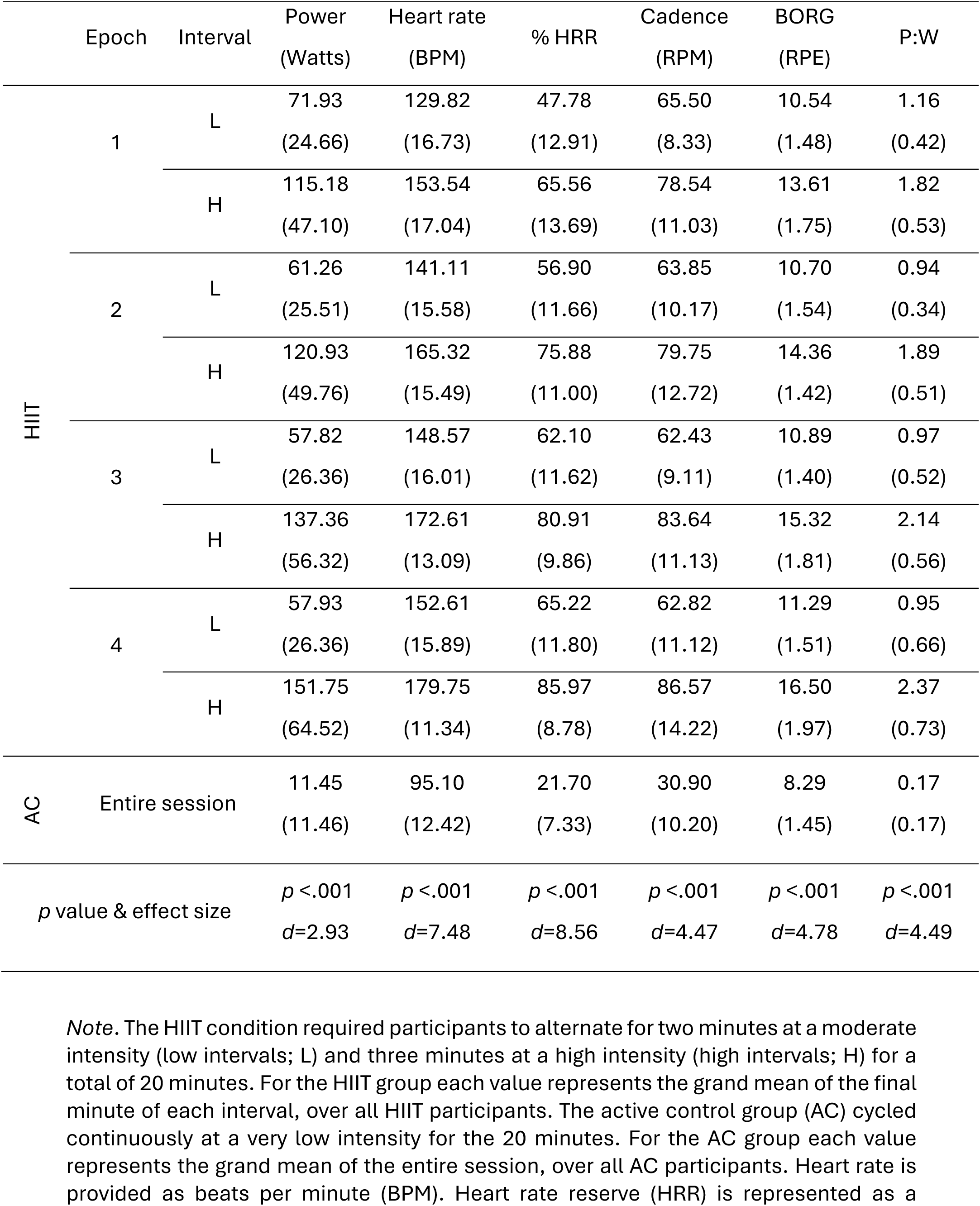
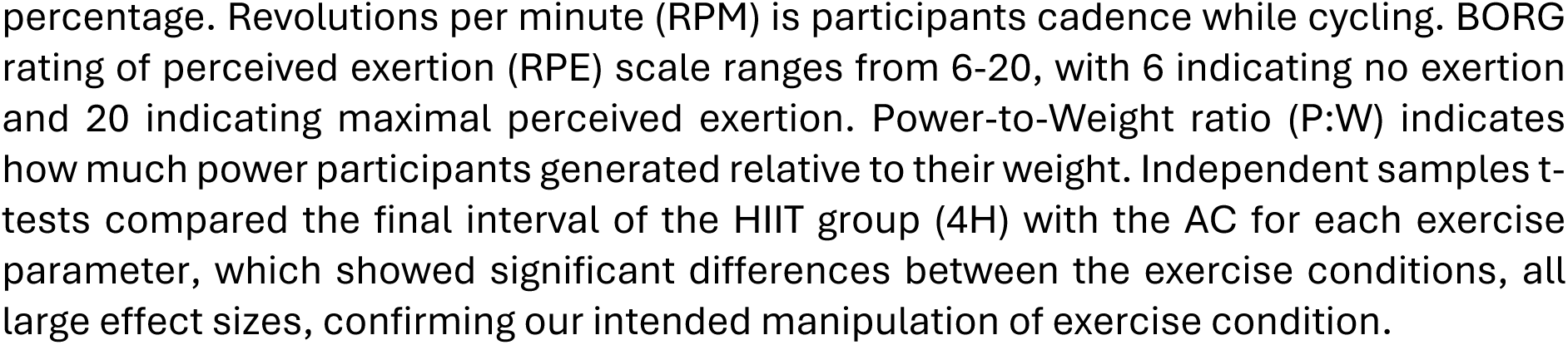
Grand mean of exercise session (standard deviations)

### Skill learning

We first examined interactions between exercise and skill in a linear mixed model. We observed a main effect of Block (*F*_5, 245_ = 11.34, *p* < .001), an interaction between Exercise Group and Block (*F*_5, 245_ = 2.37, *p* = .040), with no main effect of Exercise Group (*F*_1,49_ = 2.78, *p* = .10) Visual examination of the standardised skill plot over the session (Figure 1d) shows a steeper learning trajectory in the HIIT group from commencement of motor practice. Post-hoc contrasts revealed that the Groups differed from Block 4 of the SVIPT task (marginal mean difference between HIIT and control groups, Block 4 = 1.428, 95% CI [0.092, 2.763], *p* = .036; Block 5 = 1.460, 95% CI [0.124, 2.796], *p* = .032; Block 6 = 1.442, 95% CI [0.106, 2.778], *p* = .034).

We further calculated a single total learning metric for each participant (see methods) and confirmed greater total learning for the HIIT group (*M* = 3.07, *SD* = 1.86) compared to the active control group (*M* = 2.08, *SD* = 1.52), *t*(49) = −2.08, *p* = .043, with a medium effect size, *d* = −0.58 (Figure 1e).

### Online and offline contributions

Next, we investigated whether learning on the SVIPT task was attributable to online or offline processes (one-sample t-tests against zero, with data collapsed across groups), and secondly whether the subcomponents of total learning were differentially impacted by exercise (independent-samples t-tests). The one-sample t-tests (Figure 2a) indicated online skill improvement within blocks (*M* = 3.39, *SD* = 4.92), *t*(50) = 4.92, *p* < .001, *d* = 0.69, with no offline decrement between blocks (*M* = −0.79, *SD* = 4.56) *t*(50) = −1.23, *p* = .224, *d* = −0.17.

**Figure 2.**
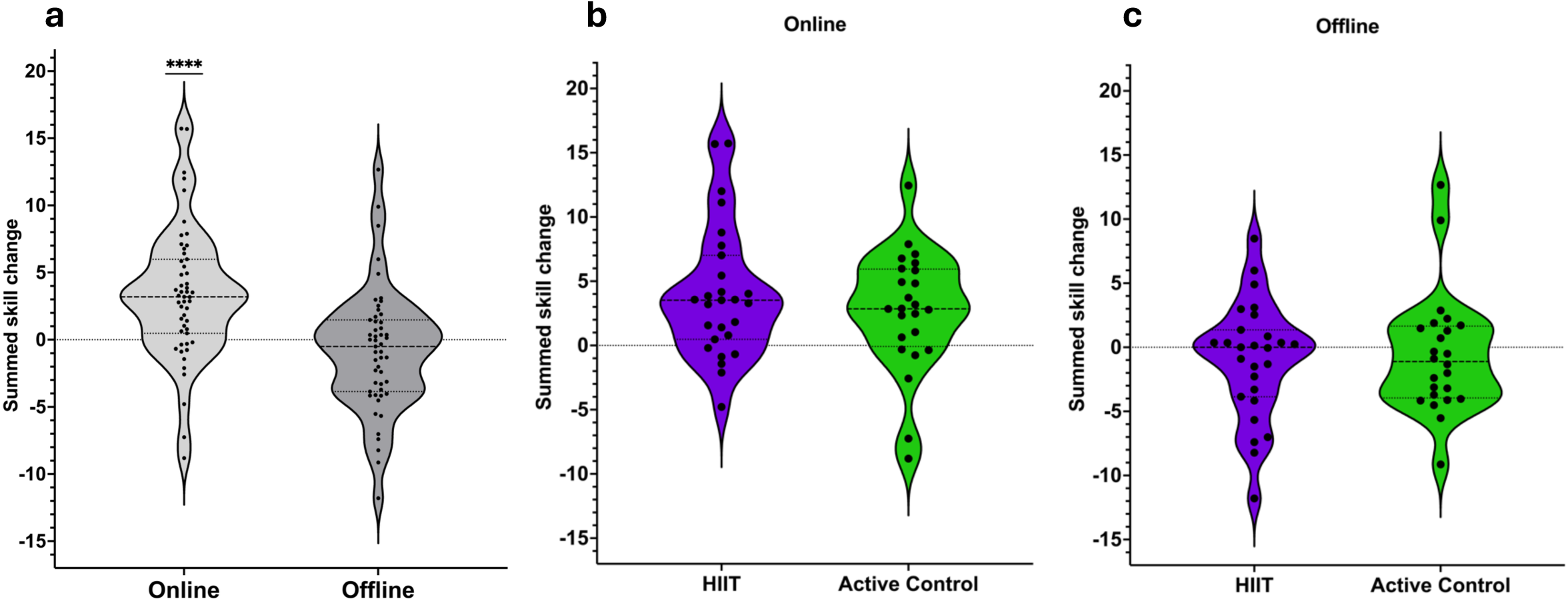
Influence of exercise on components of skill learning. *Note.* **a)** Online and offline skill changes for all participants. The skill gains on the SVIPT are primarily attributable to online active practice (light grey) (\*\*\*\**p* < .001), rather than offline periods of rest (dark grey). Comparing between exercise groups, HIIT had no significant direct effect on online **(b)** or offline **(c)** processes.

There were no significant differences between groups for either the online periods (HIIT: *M* = 4.03, *SD* = 5.14; active control: *M* = 2.67, *SD* = 4.66), *t*(49) = −0.99, *p* = .330, *d* = −0.28, or the offline periods (HIIT: *M* = −0.96, *SD* = 4.52; active control: *M* = −0.59, *SD* = 4.69), *t*(49) = 0.28, *p* = .780, *d* = 0.08 (Figure 2b&c).

### Secondary measures of skill

To better understand the interaction between exercise group and block for skill learning, we entered speed and accuracy components into linear mixed models. Across all participants, speed (trial time, reaction time, and movement time) and accuracy (force error) improved across blocks (main effect of Block *p* > .007 for all variables), indicating that across practice, participants became faster and more accurate (Figure 3). Comparing between HIIT and control groups, there were no main effects of Group for any of these additional skill components (all *p* > .05). However, an interaction of Exercise Group x Block was observed for force error (F_5, 254_ = 2.52, *p* = .03), indicating that over the session participants in the HIIT group demonstrated a greater improvement in accuracy compared to the control group (Figure 3b). Overall, these results indicate that the effect of HIIT on learning occurs via a combination of enhanced speed and accuracy, but is particularly influenced by improved accuracy.

**Figure 3.**
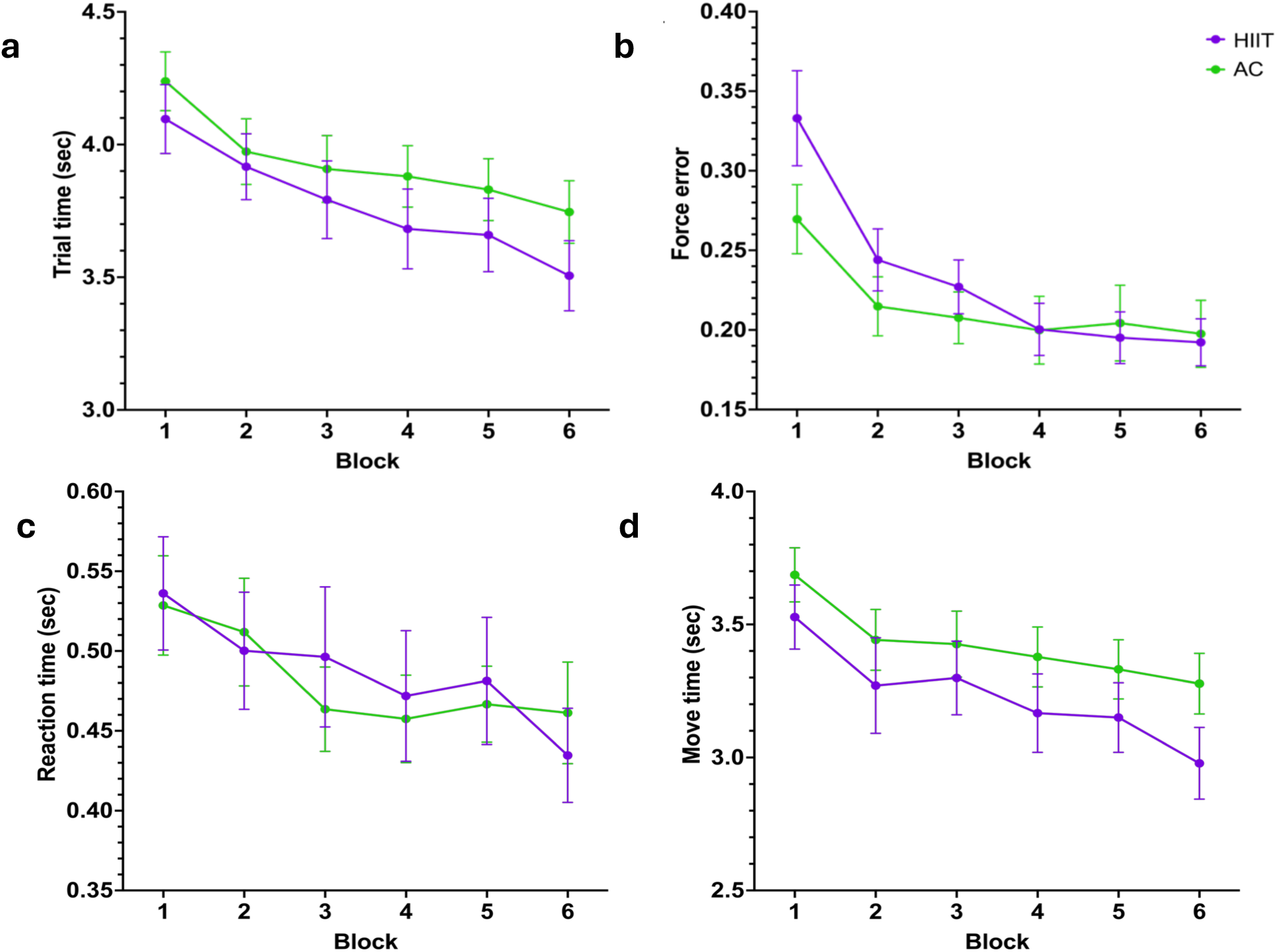
Trial time, force error, reaction time, and move time plotted across the session. *Note.* No differences were seen between HIIT and active control groups on SVIPT trial time (**a**), accuracy, represented as force error (**b**), reaction time (**c**), or move time (trial time minus reaction time) (**d**). However, an Exercise Group by Block interaction was found for force error, indicating improved SVIPT accuracy following HIIT exercise (*p* = .03).

### Genetic results

Lastly, we examined the influence of genetic polymorphisms on learning outcomes. There were no main effects of BDNF polymorphism (*F*_1, 43_ = 1.179, *p* = .28), or Exercise Group × BDNF interaction (*F*_1, 43_ = 1.579, *p* = .22) on total skill learning. Similarly, no main effects were found for DRD2/ANKK1 polymorphism (*F*_1, 45_ = 0.02, *p* = .89), or interaction effects between Exercise Group × DRD2/ANKK1 (*F*_1, 45_ = 0.74, *p* = .39) on total skill learning. No significant effects were observed when the online or offline learning measures were entered as the dependent variable (*all p* > .45).

## Discussion

In this study, we investigated the effect of a preceding bout of HIIT exercise on the acquisition of a novel motor skill, and whether improvements in skill attributable to online training or the brief offline rest periods between practice blocks. In support of the primary hypothesis, HIIT exercise enhanced skill acquisition, relative to an active control condition. However, there was no selective effect of HIIT exercise on online or offline learning processes. Additionally, there was no interaction between BDNF or DRD2/ANKK1 genotypes and the improvement in motor skill following exercise. Overall, these results add to a growing evidence base indicating a role of HIIT exercise in priming motor learning, and in particular indicate facilitation during early skill acquisition phase.

In line with the primary hypothesis, we report evidence that a preceding bout of HIIT exercise enhanced motor skill acquisition. Qualitatively an additional benefit of HIIT on skill acquisition was observed during initial blocks of practice - a trend which reached statistical significance from Block 4 onwards (i.e., ∼16.5 minutes post exercise cessation). While there is mixed evidence regarding the effect of exercise on skill acquisition^8–10,12,13,17,18^, these results are consistent with previous work demonstrating a benefit^8,12,17^. Despite highly similar methodological components such as sample characteristics, exercise protocol, and motor tasks, the mixed findings across studies to date highlight the complex interplay between these factors, as well as individual participant differences which may combine to influence the response following exercise^35^. Our results also indicate that skill improvements in the HIIT group are driven by a combination of improved speed and accuracy, but primarily accuracy. This enhanced accuracy following HIIT is consistent with previous work looking at moderate exercise and skill learning in the same motor task^12,17^. This suggests that across varying intensity exercise, skill gains on this visuo-motor task are primarily driven by improved accuracy during the task.

There are several factors which may explain the observed benefit of exercise on skill learning. Exercise, particularly performed at higher intensities, causes a cascade of physiological effects including increased cerebral blood flow, oxygen to the brain, and circulating metabolites, neurotransmitters, and neurotrophins, which ultimately creates an environment conducive for neuroplasticity, and therefore learning and memory^28,36^. Specifically, exercise is associated with upregulation of neurotrophins such as BDNF, which is known play an essential role in facilitating neuroplasticity and learning^30,37,38^, with higher intensity exercise associated with increased BDNF levels^38^. Acute exercise is also associated with increased neurotransmitters such as dopamine, serotonin, adrenaline, and acetylcholine, which are implicated in memory formation and learning^11,37,39,40^. While these substances act on various processes and timescales to support learning, specifically higher concentrations of noradrenaline and lactate following exercise have been associated with better skill acquisition^41^. Increased cerebral blood flow, cortisol, and noradrenaline following exercise are associated with improved arousal^24,37,40^, and may therefore have improved focus during task performance. In turn, these interactions may have contributed to improved accuracy, and overall acquisition of skill. Additionally, exercise is capable of modulating cortical excitability to facilitate long-term potentiation (LTP)-like plasticity to promote learning^13,20,42–44^, with higher intensity associated with greater plasticity responses^20^. Together these factors, at least in part, may explain the benefit of exercise on skill learning.

In the current study, skill improvement occurred during online, but not offline periods. These results contradict previous observations demonstrating rapid offline consolidation of motor skills for motor sequence learning tasks, such as the serial reaction time task^19,25,26^. The reason for these divergent study outcomes may relate to differences in task demands across studies. The SVIPT task has a relatively slow learning trajectory, and requires motor adaption, force control, sequence learning, and target selection linked to distributed activity across corticomotor, cerebellar, and basal ganglia networks^7,45–49^. Comparatively, the SRTT employed in previous studies has a faster learning trajectory, does not require precise force modulation, and is associated with hippocampal dynamics, such as sharp wave ripples and neural replay during rest^50–52^. Furthermore, fast vs slow learning phases are mediated by distinct neural processes and brain networks^7,46–48,53^. As such, given the different learning trajectories between the SVIPT and SRTT tasks, this may influence the relative recruitment of online and offline processes underlying within-session skill acquisition.

HIIT exercise enhanced total skill learning, but this effect was not reflected by a selective influence on online or offline periods. This suggests that the facilitation of exercise may arise through effects that impact upon both online and offline processes. This finding aligns with our previous work which observed no selective impact of HIIT on online or offline learning processes during practice of a serial reaction task^19^. These findings are interesting in the context of previous work reporting a benefit of exercise on offline processes across longer timescales (e.g., hours to days following skill training)^3,4,8–16^. Collectively, this evidence offers insights into the timescales in which cardiovascular exercise (and in particular HIIT) may influence consolidation. Indeed, via action on key neurotrophic factors, such as BDNF, exercise may maintain an environment in the brain primed for the microstructural plasticity associated with learning and memory consolidation^6,7,29,30,54,55^. Thus, in the context of consolidation, the influence of exercise may be relatively specific to retention of skill in the hours-days which follow training, rather than at a shorter timeframe, such as brief rest periods separating practice. Further research is required to establish the physiological factors mediating the time-dependent influence of exercise on motor skill consolidation.

Contrary to the final hypothesis, there was no apparent influence of BDNF or DRD2/ANKK1 polymorphisms on the facilitation of skill acquisition after HIIT exercise. While we acknowledge our relatively modest sample size in the context of genetic studies, this result aligns with previous work which found no effect of these genetic variations on initial skill acquisition following exercise^5^. This previous work however did show that DRD2/ANKK1 polymorphism impacted skill retention following exercise 24 hours after initial learning^5^. The absence of genotype effects at acquisition may reflect the timescales at which BDNF and DRD2/ANKK1 operate. BDNF-mediated neuroplasticity typically unfolds over hours to days, involving protein synthesis, dendritic remodelling, and LTP processes^29,55^ that perhaps support longer offline memory consolidation rather than immediate skill acquisition during the session. Additionally, while DRD2/ANKK1 genetic variations influence dopamine receptor density and function^56^, the acute dopamine response following HIIT exercise might be sufficient to saturate available receptors regardless of genetic variation, and may therefore have masked any differential effects. Further, as mentioned there are numerous complex physiological responses to exercise regarding neurotransmitters and neurotrophins, which collectively facilitate neuroplasticity and learning^28,30,36,37,40^. In this sense, it may be difficult to disentangle genetic interactions within this biochemical cascade via analysis of single genotypes. On this timescale of within-session online versus offline practice effects, genetic variation in BDNF and DRD2/ANKK1 did not influence skill acquisition following exercise.

There are certain limitations to this study which should be acknowledged. As participants practice the SVIPT task with the goal of being both fast and accurate, they complete each trial in a progressively shorter amount of time. However, for consistency across participants, we presented each trial on the SVIPT for a set duration. Thus, as a function of improved skill, micro-rest periods may have been introduced during online practice (i.e., brief 1-2 second breaks between trials during latter stages of practice). While the exact timescale of rapid offline ‘micro-consolidation’ remains unclear, it is plausible that certain offline consolidation processes may have contributed to learning during these micro-rest periods between trials. By extension, these brief rests may therefore have inflated online learning effects, while masking offline gains. However, as the change in speed would be most prominent in the later practice blocks, yet we observed trends in learning emerging from the beginning of practice, this timing component is unlikely to have significantly influenced our results. Additionally, considering we found task accuracy to be the prominent driver of improved performance rather than speed, the change in speed across the task is unlikely to cause significant issue. However, future research may investigate the online and offline consolidation within sessions for other motor tasks that enable more discrete delineation of online and offline periods. Additionally, further work is required to assess how intrinsic differences in motor skill/function may influence the response to exercise priming.

Overall, a single bout of HIIT exercise enhanced motor skill acquisition. Skill gains on the SVIPT were attributed to the online periods of practice, suggesting that complex visuomotor skill learning relies on online processing. There was no interaction between BDNF or dopamine D2 receptor polymorphisms and skill gains following exercise, suggesting the mechanisms of these factors may not operate on this within-session timescale.

## Methods

### Experimental Design

This study utilised a mixed factorial design [between-subject factor of Exercise Group, within-subject factor of SVIPT block] to investigate the effect of acute exercise on the acquisition of a novel motor skill. Participants were pseudo-randomly allocated to either a HIIT exercise condition or a time-matched low intensity, active control group, minimising between-group variance with respect to age, biological sex, and baseline physical activity level. This study was approved by the Monash University Human Research Ethics Committee (MUHREC 27742), and all participants provided informed consent prior to participating.

### Participants

The sample consisted of 51 right-handed, healthy young adults recruited from the general community (57% female), ranging in age from 18 to 38 years old (*M* = 22.04, *SD* = 3.41). To be eligible, participants were required to have no contraindications to exercise (e.g., physical injury, asthma, blood pressure problems, family history of heart disease), no history of psychiatric or neurological illness or injury, and not be currently taking psychoactive medications.

### Procedure

Participants completed a one-hour testing session involving a bout of high-intensity exercise and motor skill training. Lifestyle/physiological measures were first recorded, including, height, weight, and resting heart rate (RHR), as well as self-reported measures of handedness (Edinburgh Handedness Inventory), exercise habits (International Physical Activity Questionnaire, IPAQ), and sleep quality (Pittsburgh Sleep Quality Index, PSQI). Participants also provided a saliva sample for genetic analysis. Participants then completed the exercise and motor skill training (Figure 1a-b)

#### Exercise Protocol

Participants completed a 20 minute supervised bout of exercise on a cycle ergometer^57^. Exercise intensity was tailored to each individual, based on their heart rate reserve (HRR) which was determined by subtracting their RHR from their age predicted maximum (220 – age). Heart rate was continuously monitored during using a single lead electrocardiogram^58^ throughout the duration of each session. The HIIT protocol is well established and has been utilised in our previous work^13,18–20,32,59,60^. In brief, this protocol involves alternating epochs of two-minute high intensity (target heart rate of up to 90% HRR) and three-minute moderate intensity cycling (target heart rate of 50-60% HRR), for 20 minutes in total. This was followed by a 1–2-minute cool down on the bike to allow for heart rate recovery. For the low intensity condition, participants cycled at a very low cadence with minimal exertion (i.e., the target heart rate was monitored and kept below 20% HRR). The active control participants watched a documentary of their choosing to maintain engagement and attention throughout the low intensity exercise. Participant’s rating of perceived exertion (RPE) was recorded throughout the exercise using the original version of the BORG scale, where ratings range from 6 (no exertion) to 20 (maximal exertion)^61^.

#### Sequential visual isometric pinch task (SVIPT)

Participants were seated in front of a computer screen and asked to grip a force transducer between their index finger and thumb of their dominant hand. Squeezing the force transducer moved a cursor horizontally on the computer screen, proportional to the applied force. Five targets appeared on the screen, and participants were instructed to produce five force pulses to move the cursor to the middle of each coloured target in a specific pre-determined order (i.e., red, blue, green, yellow, white), returning to the start position between pulses. Participants were instructed to perform the task as quickly and as accurately as possible.

Prior to exercise, participants received task instructions, and their maximum voluntary pinch contraction (MVC) was measured. MVC was defined as participant’s maximal pinch force output across ∼1s period on the force transducer. Participants were given three attempts at generating an MVC, with a ten second rest in-between, and the maximum value obtained was taken as MVC. This allowed SVIPT practice to be tailored to each individual, with the furthest target on each trial requiring a force output equivalent to 45% of individual MVC to minimise fatigue. Participants then completed three practice trials with targets appearing in a simple ascending sequence, in contrast to the order presented during testing (blue, white, green, red, yellow). The practice block was administered to familiarise participants to the task and ensure they could register five force pulses within the allotted trial duration.

Critically, the task was partitioned into online practice periods consisting of six blocks of 10 trials (60 trials total), and offline rest periods of 30 seconds duration between blocks. Seven seconds were provided for participants to complete each trial, with three seconds in between each trial. In total, the task took 15 minutes to complete (12.5 minutes of ‘online’ task practice, and the 5 rest periods of 30 seconds). Motor skill training on the SVIPT^62^ was timed to commence 10-minutes after exercise, in line with previous studies reporting a maximal priming effect when there is a close temporal coupling between exercise and learning^1,20,22,23^.

### Genetic Analysis

Saliva samples were acquired using Oragene DNA self-collection kits (DNA Genotek). Genotyping was completed at the end of all data collection to reduce possible bias. DNA was isolated using standard procedures recommended by the supplier. The BDNF val66met and DRD2/ANKK1 glu713lys polymorphisms were genotyped using TaqMan SNP genotyping assays (rs6265 and rs1800497) on a Light Cycler 480 system (Roche).

### Data Analysis

Data analysis of the SVIPT task was conducted in MATLAB (version R2023b), and statistical analysis was performed in JASP (version 0.19.1). For each SVIPT trial, speed and accuracy were calculated (Figure 1c). Trial speed was defined as the time from initial target onset to the final force pulse execution on each trial. Accuracy was calculated as an error score from the summed differences between each of the five individual force peaks (i.e., force outputs to move the cursor to each of the five targets) relative to the centre point of each target. Individual trials were excluded if the participant produced more or less than five force peaks, or failed to make an adequate attempt to return to the start/home position between pulses. Out of the 3060 trials conducted, 140 were excluded, i.e., 4.58% of all trials (per participant *M* = 2.75%, *SD* = 2.56 of trials).

In line with previous studies that have employed the SVIPT^13,27,62^, motor skill was calculated according to a function that includes both speed and accuracy as inputs. To determine this function, a separate data set (*n* = 10) was acquired to determine the speed-accuracy trade-off function when performing the SVIPT task^63^. In brief, this independent sample of healthy adults was required to complete the task at different speeds paced by a metronome (10 blocks of eight trials across two separate days). Performance was then averaged, and a speed-accuracy trade-off function was fit to the data:

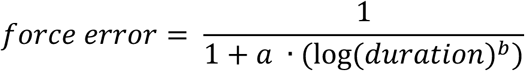

where ‘duration’ is the mean trial time of each block, and a and b are dimensionless parameters. The resultant function (r^2^ = .99) derived the value of parameter b as 1.627. Rearranging this function allows for determination of the skill parameter (‘a’), whereby:

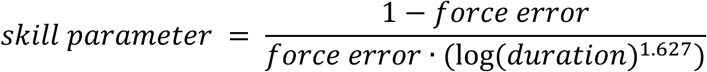

Higher values of the skill measure reflect performance that is both fast and accurate (i.e., a translation of the entire speed-accuracy trade-off function to the left).

To model learning trajectories across the session, participants’ skill measure was standardised so that scores represent a ratio of their performance relative to block 1, with increasing values representing improved performance. A linear mixed model analysis was conducted to examine skill learning across blocks, and between exercise groups. Group, and block were entered into the model as fixed effects, with participant included as a random effect (S*kill ∼ Group x Block + (1 | Participant))*. Planned contrasts were then used to identify when group differences began to emerge.

To examine the breakdown of learning, each participant’s raw skill values were averaged over the first and last three trials of each block to determine skill at the start and end of blocks respectively. Total learning was calculated as the change in skill across the session (end of block 6 - start of block 1). Change in skill was then further partitioned into “online effects”, the summed change in skill within blocks (end of block n – start of block n), and “offline effects”, the summed change in skill between blocks (start of block n+1 – end of block n). As per our previous work^19^ one-sample t-tests were used to determine the significance of online and offline effects over the entire sample, and three independent samples t-tests were used to compare total learning, online and offline effects between the HIIT and active control groups.

Trial time, force error, reaction time, and move time (trial time minus reaction time) were also analysed to assess which components of skill may be influenced any observed effects. A linear mixed model with Exercise Group, and Block entered into the model as fixed effects, and participant as a random effect, was run for each dependent variable (trial time, force error, reaction time, and move time). An alpha level of .05 was set for all analyses.

#### Genetics analysis

Genotypes were dichotomised as common homozygotes (Val/Val for BDNF, and Glu/Glu for DRD2/ANKK1), or carriers of the polymorphism allele (Met for BDNF, and Lys for DRD2/ANKK1), which included heterozygotes (Val/Met, and Glu/Lys) and rare homozygotes (Met/Met, and Lys/Lys) to account for differences in population frequency of occurrence. Genotype dichotomy for BDNF and DRD2/ANKK1 were entered into two separate 2×2 ANOVAs to investigate the effects of Genotype and Exercise Group on total skill learning over the session.

## Data availability

De-identified behavioural data are available at [https://osf.io/ae4hr/]. All code used for analysis is available at [https://github.com/jhendrikse/].

## Acknowledgements

This study was funded by an Australian Research Council Discovery Grant (DP200100234; J.C, W.B, and Z.H) and Future Fellowship awarded to J.C (FT230100656). J.H is supported by an Australian Research Council Discovery Early Career Research Award (DE240101348). The funder played no role in study design, data collection, analysis and interpretation of data, or the writing of this manuscript. Aristidis-Peter Lavouasier, Liam Stanborough, Lachlan Terry contributed to data collection.

## Author information

### Author contributions

J.H and J.C supervised the project. E.B, J.K, J.H, N.A, J.H, and J.C designed, conducted data collection, and analysed the data for this project. E.B, J.K, J.H, and Z.H conducted genetic analysis. E.B, J.H and J.C wrote the manuscript. All authors read and approved the final version of the manuscript.

Corresponding author:

Correspondence to either Dr. Joshua Hendrikse or A/Prof. James Coxon

